# Global patterns and predictors of PFAS contamination in odontocetes

**DOI:** 10.64898/2026.03.04.709656

**Authors:** Lavinia Stokes, Karen A. Stockin, Gavin Stevenson, Jesuina de Araujo, Frédérik Saltré, Katharina J. Peters

**Author notes:** Correspondence **Katharina Peters****, Lavinia Stokes**.

## Abstract

Per- and polyfluoroalkyl substances (PFAS) are globally recognised as emerging contaminants of concern due to their persistence, toxicity, endocrine-disrupting and immunosuppressive effects. Because of their extensive industrial use, PFAS are now widespread across ecosystems and accumulate in marine environments. Despite their ubiquity, the extent and drivers of PFAS contamination remain poorly characterised, particularly in marine systems. Odontocetes (toothed whales) are effective bioindicators of marine pollution, integrating contamination across regions, time, and trophic levels.

Here, we present the first global assessment of factors influencing PFAS contamination in marine ecosystems by analysing standardised PFAS concentrations of PFNA, PFDA, PFUnDA, PFDoDA and PFOS reported for 713 liver samples across 33 odontocete species spanning 13 countries from 2000 to 2023. Using generalised linear mixed models, we evaluated the effects of genus, location, sex, life stage, and sampling year on PFAS concentrations, combining published datasets with new samples from Australia. Genus and location were the strongest predictors, suggesting that interspecific ecological and physiological traits likely contribute to PFAS accumulation. Concentrations were highest in males and younger individuals, consistent with maternal offloading and possible age-related dilution. Spatio-temporal trends indicate that PFAS contamination is widespread and increasing globally, with highest concentrations reported in the Pacific.

This study provides a critical baseline for understanding global PFAS exposure in marine mammals, which underscores the need for coordinated monitoring and further research to address regional data gaps and potential unrecognised biological effects.

**Highlights:**

- High genus-specific and spatial differences in PFAS contamination across odontocetes globally.
- Increased contamination in younger/smaller individuals.
- Sex-specific trends, including higher PFAS levels in male odontocetes.
- Spatio-temporal trends suggesting increased PFAS concentration despite global regulatory efforts, with highest concentrations in the Pacific Ocean.

## 1. Introduction

The unique molecular properties of per- and polyfluoroalkyl substances (PFAS) have led to their widespread application in almost all modern industrial processes and consumer products (e.g., water-repellent clothing, cookware with non-stick coating), with more than 1,400 PFAS applied across 64 categories, spanning over 200 applications (Abunada et al., 2020; Gaines, 2023; Glüge et al., 2020). However, PFAS pose considerable risks to both human and animal health (Abunada et al., 2020; Fair & Houde, 2018), including, neurological (Cao & Ng, 2021; Pedersen et al., 2015), immunological (Beans, 2021; Ehrlich et al., 2023; Fair et al., 2013), and reproductive effects (Custer et al., 2012; Hoffman et al., 2010; Saikat et al., 2013; Vélez et al., 2015; Wee & Aris, 2023; Wood et al., 2021). Increasingly, PFAS are considered ‘contaminants of emerging concern’, which describe potentially deleterious chemicals where there is insufficient understanding on its presence and effects in the environment (Feng et al., 2023; Schnoor, 2003).

The extensive use of PFAS inevitably results in its widespread dispersion into the marine environment, primarily from direct sources, including manufacturing effluent (Pétré et al., 2022), wastewater facilities (Lenka et al., 2021) and military training bases (Ruyle et al., 2023; Skutlarek et al., 2006). Therefore, higher concentrations of PFAS are more likely in locations closer to anthropogenic activities (Ahrens & Bundschuh, 2014; Podder et al., 2021) and the concentration and variability of contamination is expected to reflect regional pollution levels (Tansel, 2024). Additionally, PFAS are released into the environment through many diffuse, indirect pathways including industrial and agricultural runoff (Peter & Lee, 2025), sewerage (Ford & Ginley, 2024) and atmospheric deposition (Ahrens et al., 2011; Jarvis et al., 2021; Podder et al., 2021). Temporal and spatial variability of PFAS contaminants are likely driven by ocean dynamics such as frontal processes, upwelling, and mixing, influencing the dispersal and concentration of contaminants (Cai et al., 2018; Lohmann & Belkin, 2014; Schlosser et al., 1995; Wang, 2018). Furthermore, regulatory efforts likely reflect variability of PFAS contamination such as decadal decreases in PFAS contamination levels in the Baltic Sea observed since PFAS phaseouts in the region since 2002 (Soerensen et al., 2024).

Marine biota are highly susceptible to PFAS contamination and useful bioindicators for assessing environmental contamination through space and time (Bossart, 2011; Plön et al., 2024) as marine ecosystems serve as reservoirs or the ‘ultimate sink’ for these emerging contaminants of concern (Bashir et al., 2020). Once introduced into marine waters, PFAS can dissolve, form solutions, or attach to suspended particles (Eisma, 2012), entering marine biota via the water column and food intake (Suedel et al., 1994). Once ingested, PFAS may amplify within organisms through bioaccumulation (Bryan & Darracott, 1979; Mackay & Fraser, 2000) and/or biomagnification processes (Gray, 2002). Marine mammals are reported to have, along with avian species, the highest PFAS biomagnification potential among wildlife (Kelly et al., 2009; Tomy et al., 2009) likely due to their diet, respiratory exchange mechanisms (De Silva et al., 2021) and their longevity.

Odontocetes (toothed whales) are particularly useful for assessing and monitoring PFAS in our oceans given their widespread global occurrence, comprising more than 70 species across marine and some freshwater habitats (Hooker, 2009), and because they occupy meso- and apex predator positions in food webs. These characteristics provide a unique opportunity to examine differences in PFAS contamination in relation to their diverse ecology and physiology. Spatial exposure is a major driver of PFAS concentration in odontocetes (Foord et al., 2024; Galatius et al., 2013) as species inhabiting coastal regions and/or areas with high levels of PFAS production, use, and historically inconsistent regulation (Buck et al., 2011; Tansel, 2024) are likely to experience higher contamination. This elevated exposure due to the proximity to coastal runoff and anthropogenic processes (e.g., wastewater discharge, airports, fuel refineries in ports and military training bases, (Anderson et al., 2023; Hu et al., 2016; Kanaji et al., 2017; O’Shea, 1999). Thus, biomonitoring has become important for assessing overall ecosystem contamination in marine environments (Gewurtz et al., 2011; Oakes et al., 2010). While some PFAS compounds are currently regulated, many new PFAS are entering the market, resulting in mixed and sometimes contradictory trends in temporal contamination studies (Bernardini et al., 2021; Law, 2014). At present, establishing robust links between management actions and PFAS contamination in odontocetes, including clear global temporal trends, remains challenging.

Beyond environmental and spatial exposure, species-specific biological characteristics of odontocetes, such as diet, sex and life stages might also influence PFAS accumulation (Fair & Houde, 2018; Muir & Miaz, 2021; Powley et al., 2008). Diet can influence biomagnification effects, where contamination of PFAS traverses lower-trophic prey resulting in highest concentrations in the tissue of upper-level predators (Gray, 2002).

Comparing PFAS contamination amongst sexes, males generally exhibit higher contaminant concentrations compared to females across animal species worldwide (Bangma et al., 2017), likely due to differences in physiological processes (Lynch et al., 2019) and maternal transfer mechanisms (Dorneles et al., 2008; Kelly et al., 2009; Lemos et al., 2024), resulting in mature females offloading PFAS contaminants to their calves (Gaylard, 2017; Lemos et al., 2024).

Additionally, age and body size may further be linked to the process of bioaccumulation, where older and thus larger individuals may display higher contaminant levels, as observed for wider contaminants in odontocetes (Lavery et al., 2008; Machovsky-Capuska et al., 2020). Differences in properties and behaviour of PFAS compounds (i.e., sulfonic acids such as PFOS, PFBS, PFHxS, carbonic acids such as PFOA, PFNA, PFDA) within animals, including elimination, half-life, or transformation ability, can also impact the persistence and bioaccumulation (Alalm & Boffito, 2022; Semerád et al., 2022), highlighting the importance of assessing these different chemical classes separately.

While PFAS contamination in odontocetes has been analysed in different locations worldwide (recently reviewed in Foord et al., 2024), most studies focused only on one or a few species and locations. Yet there has been no quantitative global assessment comparing PFAS contamination across odontocete samples, thus hindering current understanding of global large-scale driving patterns of PFAS. Here, we evaluate the impact of genus, time, location, sex and life stage on PFAS concentration in the marine environment worldwide, using odontocetes as bio-indicators. Specifically, we use generalized linear mixed models on a comprehensive global dataset of PFAS concentrations in odontocetes to test the following hypotheses: (1) PFAS contamination differs across genus groups, with coastal species recording higher concentrations than counterparts occurring offshore; (2) PFAS concentration is higher in regions where PFAS are primarily manufactured or less regulated, compared to areas with limited production and stricter regulatory controls; (3) PFAS concentration increases through time due to the increase in manufacturing and use, and; (4) PFAS concentration is higher in males than in females, as females potentially eliminate PFAS via maternal offloading.

## 2. Methods and materials

### 2.1 Global literature search and database compilation

We compiled data on PFAS concentrations in odontocetes from peer reviewed (journal articles) and grey literature (government reports, theses) retrieved from Google Scholar, Scopus, Science Direct and PubMed. Search keywords included ‘PFAS’, ‘perfluoroalkyl’, ‘polyfluoroalkyl’, ‘contaminant’, ‘dolphin’, ‘whale’, ‘porpoise’, ‘odontocete’ (used in different combinations). To limit the selection to only papers relevant to this study, we established a series of inclusion criteria: *i*) date restriction: only data analysed from the year 2000 onwards were included. This cut-off was chosen to address advancements in PFAS detection methods, as older studies often had limited capacity to detect a broad spectrum of PFAS compounds due to technological constraints; *ii*) species restriction: the study species had to be an odontocete to control for diet, metabolic processes, and other large-scale ecological differences that mysticetes may have; *iii*) tissue-specific analysis: only PFAS analysis of liver was included to account for heterogeneity in PFAS burden between tissues in contaminant storing and metabolic processes.

We compiled the data that passed these first set of criteria into a global database and then applied additional secondary criteria to account for variations in detail and reporting styles across studies. To remain in the database, papers needed to provide: *iv*) the country that the sample had been collected from; *v*) total body length of the specimen. Total body length was cross-checked with known total body lengths for each species and sex in the literature (Table S10), including minimum recorded calf size for each species to exclude foetuses; *vi*) the sex of the specimen; *vii*) the year of death of the animal. For samples that did not have an accurate date of death, the later year was assumed (e.g., 2022/2023, 2023 was assumed); *viii*) individual PFAS compound contamination: results had to be provided for every PFAS compound detected, rather than just the total sum. Only wet weight (w/w) concentrations were recorded in the database. We used ng/g, transforming other units (e.g., mg/kg) to ng/g as required.

A total of 710 samples across 18 papers were included in the final analysis (Table S9). Only five compounds were tested consistently across all studies and thus used in the final analyses: four long-chain carboxylates (PFNA, PFDA, PFUnDA, PFDoDA) and the long-chain sulfonic acid PFOS (see S5). To complement the published data, we also added three new samples from southeastern NSW, Australia, to our analyses: one common dolphin (*Delphinus delphis*) and two Indo-pacific bottlenose dolphins (*Tursiops aduncus*). Details on sampling and the analytical approach are provided in S1, S2, S3 and S4. This resulted in the first global assessment of PFAS contamination in odontocetes, analysing 713 samples from 33 species worldwide.

### 2.2 Predictor variables

We considered five predictors in our dataset which included two continuous variables (1) year of death, to account for potential changes in PFAS concentrations over time, including changes in manufacturing, usage and management and (2) life stage (measured via the body length index, see 2.3), to control for biological effects of growth and maturity on PFAS accumulation, including bioaccumulation, biomagnification, and the potential for biotransformation abilities across different life stages. A further three categorical variables included (3) genus group (where we categorised 33 species into 21 genera (Table 1, Table S12, Table S9), noting some genera compromised a single species due to limited representation in the available data) and (4) location (six categories; Pacific, Oceania, North Atlantic, South Atlantic, Arctic, Mediterranean Sea, Table S9) based on their shared oceanic basins and environmental relevance regarding PFAS pollution. Simplifying the geographical and species categories helped maintain statistical power by reducing the number of categories in the model, thus avoiding multicollinearity issues and model overfitting, while still accounting for unique ecological characteristics and physiological differences across species groups and locations. Additionally, (5) sex (two levels, male and female), was included to assess potential sex-specific differences influencing PFAS accumulation. Females were used as the reference group, which serves as the reference category for comparison to other groups, in this case males, as the literature suggests males are more likely to exhibit higher PFAS concentrations (Fair & Houde, 2018; Han et al., 2012; Lynch et al., 2019). We also tested the interaction between sex and life stage, as the relationship between life stage and PFAS concentration levels may differ between males and females due to maternal transfer.

**Table 1.**
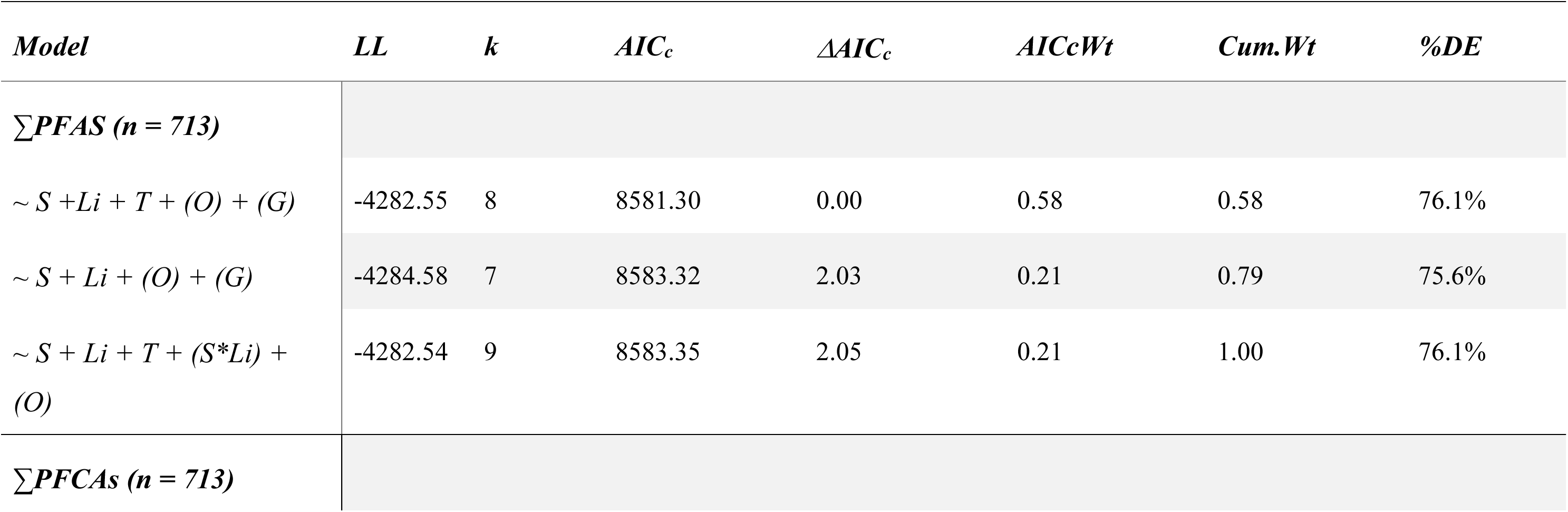
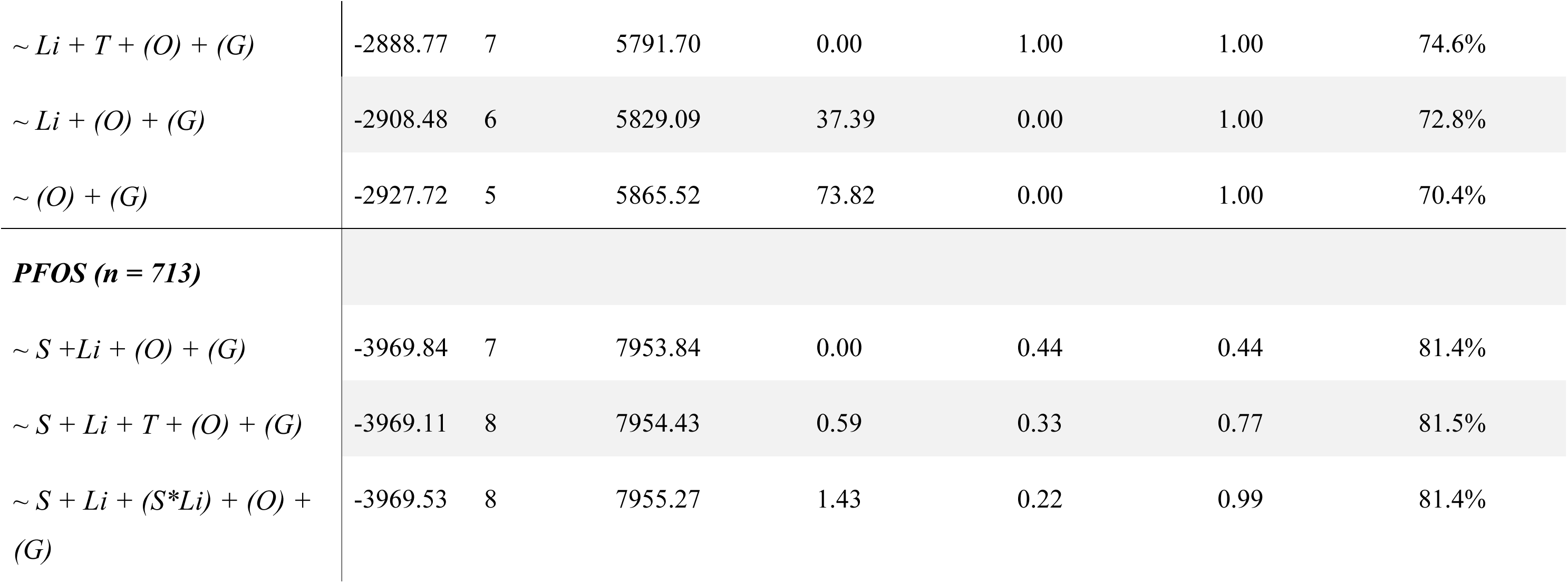
Model comparison results from generalised linear mixed models (GLMMs) predicting log-transformed ∑PFAS concentrations in odontocetes (n = 713), ranked by AIC_c_. Shown are the model structure, maximum log-likelihood (LL), number of parameters (k), corrected Akaike’s information criterion (AIC_c_), change in AIC_c_ (ΔAIC_c_), AIC_c_ weight (AIC_c_Wt), cumulative weight (Cum.Wt), and percentage deviance explained (%DE). Fixed effects include sex (S), stage of life index (Li), and year (T); random effects include Ocean (O) and genus (G). Only models performing better than the base model are shown.

### 2.3 Body length index

Determining the age of wild odontocete species is challenging due to their long lifespans and wide spatial distribution (Read et al., 2018). Accurate age estimates require costly techniques such as long-term monitoring (Connor & Krützen, 2015), tooth sample analysis (Hohn et al., 1989; Palmer et al., 2023), or epigenetic techniques like DNA methylation (Hanninger et al., 2025; Peters et al., 2023). For these reasons, age is often prohibitive in toxicology reports, as we found across studies on PFAS contamination in odontocetes. However, most studies provide morphological measurements upon necropsy, thus, here we used total body length as an indicative proxy to estimate the stage of life. To standardise for species-specific size differences, we calculated a total body index following Stockin et al. (2025) using currently known minimum calf total body length, maximum female and maximum male total body length for each species (Table S10). For each sex and species, we scaled body length data for the samples between 0 and 1, where 0 corresponds to the smallest and thus youngest individual, and 1 represents the largest and thus oldest animal ever recorded. The resulting body length index provides an indication of the animals’ stage of life and allowed comparisons across species.

### 2.4 Statistical analysis

Prior to model fitting, we quantified the proportion of variance attributable to the hierarchical structure of the data, specifically genus (taxonomic group) and location (ocean), by calculating the intraclass correlation coefficient for each grouping factor. The intraclass correlation coefficient represents the fraction of total variance in *total PFAS concentrations* explained by between-group variability rather than within-group differences. We also computed the design effect to assess the degree to which variance estimates are inflated due to clustering within these groups. Based on these results, we included both genus and location as random effects in our models (see Model justification in S11). To identify which predictors (i.e., genus, location, year, life stage, sex) best explain the variation in PFAS contaminant concentration patterns in odontocetes, we built and compared generalized linear mixed-effect models (GLMMs) using the *glmmTMB* package (McGillycuddy et al., 2025) in R version 4.4.1 (Team, 2025). We tested three metrics characterising PFAS concentration (ng/g) as response variables: (*i*) the sum of five PFAS compounds (∑PFAS: PFNA, PFDA, PFUnDA, PFDoDA and PFOS), (*ii*) the sum of long-chain PFCAs, (∑PFCAs: PFNA, PFDA, PFUnDA, PFDoDA), and (*iii*) the PFSA compound, **‘**PFOS’. We log-transformed all response variables to reduce skewness and stabilise the variance. We then fitted the GLMMs using a Tweedie distribution with a log-link, which accommodates non-normal residuals while modelling the mean-variance relationship of the Tweedie family.

We fitted six models for ∑PFAS and PFOS response variables, and five models for ∑PFCAs due to differences in predictors. The models included genus and location as random effects, corresponding to all combinations of fixed effects and the interaction between sex and life stage. For each model, we computed the Akaike’s Information Criterion corrected for small sample sizes (AIC_c_) and, following the principle of parsimony, selected the top-ranked model as the one with the lowest AIC_c_. We also computed the evidence ratios to quantify relative model supports. Model diagnostics included checks for overdispersion, residual patterns against fitted values, influential observations, and the adequacy of the Tweedie distribution variance structure. We also evaluated the relevance of the log-link function using partial dependence plots for evidence of nonlinearity.

## Results

### 3.1 Data collection

We screened >1,000 publications via the literature search, resulting in 43 papers with >1,300 samples meeting the initial set of inclusion criteria. However, only 18 of these studies provided sufficient information to satisfy the secondary criteria. Furthermore, at least 57 individual PFAS compounds or groups were identified initially across studies in odontocetes. However, only 5 of these compounds (PFNA, PFDA, PFUnDA, PFDoDA, PFOS) were consistently selected for analysis across all studies and thus were included in our analysis.

Data were further supplemented by three samples collected from southeastern Australia, resulting in a final dataset of 713 samples spanning 33 species, across 13 countries (Table S9). Detailed results of the three new samples are given in S6 and S7 and S8.

### 3.2 Factors associated with ∑PFAS contamination

The best-fitting model for total ∑PFAS concentration in odontocetes included life stage, sex and year, with genus and location as random effects (Table 1, S15; evidence ratio = 2.78 relative to the second-best model). Including the interaction between life stage and sex did not improve model fit. The best-fitting model explained 76.1% of the total variance (Table 1) with fixed effects that accounted for 1.4% (Table 1). According to the model, ∑PFAS concentration decreased by approximately 15% with increasing stage of life (exp(0.162) = 0.85, Table 2, Fig. 2). Additionally, ∑PFAS concentration were ∼ 29% higher in males than in females (exp(0.251) = 1.29) and increased by ∼14% over time (exp(0.129) = 1.14).

**Table 2.** Generalised linear mixed model (GLMM) results predicting log-transformed ∑PFAS concentrations in odontocetes (n = 713). Estimates, standard errors (SE), test statistics, and p-values are shown for fixed predictor variables: sex, age, and year. Variance (σ) and standard deviation (SD) are reported for random intercepts associated with Location and Genus group.

| **Fixed effects** | **Estimate** | **SE** | **Statistic** | **p-value** |
| --- | --- | --- | --- | --- |
| (Intercept) | 3.66 | 0.48 | 7.59 | <0.0001 |
| Sex | 0.25 | 0.10 | 2.43 | <0.01 |
| Age | -0.16 | 0.043 | -3.81 | <0.0001 |
| Year | 0.13 | 0.063 | 2.02 | <0.01 |
| **Random effects** | **$\sigma$** | **SD** |  |  |
| Ocean | 0.52 | 0.72 |  |  |
| Genus | 2.17 | 1.47 |  |  |

Random effects had the strongest effects on ∑PFAS concentrations. For the genus effect, ∑PFAS concentrations were higher than the mean in *Sousa* (+18×), *Tursiops* (+17×), and *Neophocoena* (+9×, Fig. 1). Conversely, *Lagenorhynchus*, *Pseurdorca* and *Orcinus* showed the largest negative deviations with levels between 70-80% below expected values. Among locations, the Pacific and Oceania showed the highest positive deviations from the global mean, followed by the North Atlantic (+1.28×). Conversely, the Mediterranean Sea showed concentrations ∼34% below the baseline, while the Arctic Ocean and South Atlantic were near or slightly below the global average (Fig. 1).

**Figure 1.**
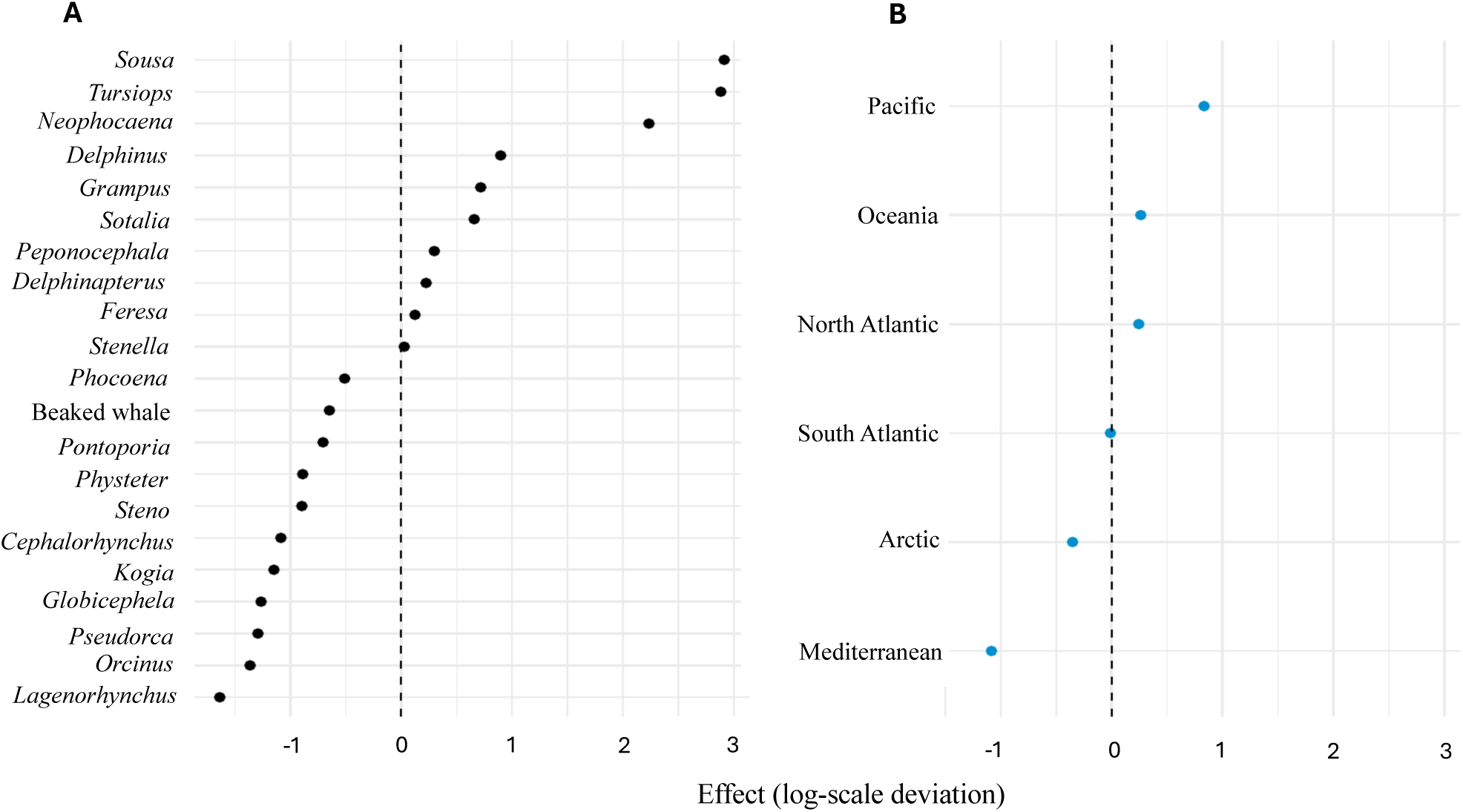
Variation in random intercept effects for genus groups A (black) and ocean locations B (blue) from the model predicting log ∑PFAS concentrations. Values represent deviations from the overall intercept on the log scale, with positive values indicating higher contamination and negative values indicating lower contamination relative to the model’s grand mean.

### 3.3 Factors associated with ∑PFCAs and PFOS contamination

The best-fitting model for ∑PFCAs concentration included life stage and year as fixed effects, with genus and location as random effects (Table 1), with an evidence ratio of 1.31 ×10^8^ relative to the second-best model. This model explained 74.6% of the total variance (Table 1) with fixed effects that accounted for 5.7% and showing that ∑PFCAs concentrations decreased by approximately 17.1% with increasing life stage (exp(−0.18808)) and increased by ∼44.9% over time (exp(0.37070), Table S13). Random effects contributed most to the variation in ∑PFCAs concentrations among genera, *Tursiops*, *Neophocaena*, and *Sousa* which showed the highest positive deviations from the mean, with levels ∼3×, ∼5×, and ∼6× higher, respectively. The lowest concentrations were found in *Pontoporia* (∼71% below the mean), *Lagenorhynchus* (∼82% below), and *Cephalorhynchus* (∼83% below). Across oceanic regions, the Pacific showed the highest ∑PFCAs levels (∼3× the mean), followed by the North Atlantic (∼1.5×) and the Mediterranean Sea (∼1.4×). In contrast, the South Atlantic and Oceania showed markedly lower concentrations, at ∼50% and ∼67% below the global average, respectively.

The best-fitting model for PFOS concentration in odontocetes included *life stage* and *sex* as fixed effects, with *genus* and *location* as random effects (Table 1, evidence ratio = 1.52 relative to the next best model), explaining 81.4% of the total variance. Fixed effects accounted for 0.7% of the total variance and showed that PFOS burden decreased by ∼13.3% with increasing *life stage* (exp(−0.14295) = 0.87) and was ∼24% higher in males compared to females (exp(0.21511) = 1.24) (Table S14). Random effects had the strongest effects on PFOS concentrations among genera, *Sousa*, *Neophocaena*, and *Tursiops* which exhibited the highest positive deviations from the mean — ∼47×, ∼22×, and ∼20×, respectively. In contrast, *Lagenorhynchus*, beaked whales, and *Pseudorca* showed concentrations ∼79–84% below expected values. Across oceanic regions, Oceania had the highest positive deviation (∼2.3× the global mean), followed by the Pacific (∼1.4×) and North Atlantic (∼1.2×). Conversely, the Arctic and Mediterranean Sea showed <u>∑</u>PFOS concentrations ∼50% and ∼60% below the global average, respectively.

## Discussion

### 4.1 Genus-specific differences on ∑PFAS contamination

Our results show that genus was the strongest predictor of total PFAS contamination in odontocetes comparative to all other variables tested (Fig. 1), which is likely attributed to ecological and physiological differences between species (Fair & Houde, 2018; Lemos et al., 2024; Muir & Miaz, 2021). The importance of assessing multiple species when looking at PFAS contamination is emphasized in the marine mammal literature (Galatius et al., 2013; Gebbink et al., 2016; Moon et al., 2010; Stockin et al., 2025; Yeung et al., 2009) due to general trends observing differences in exposure rates which likely are linked to diet and environment (De Silva et al., 2021; Fair & Houde, 2018; Powley et al., 2008). For example, in Greenland PFAS concentration in polar bears (*U. maritimus*) were higher than in ringed seals (*Pusa hispida*) (Gebbink et al., 2016). Our results suggest that even species within the same suborder, in this case Odontoceti, can exhibit inter-species differences in PFAS exposure (Fig. 1, Table 2). This is further supported by varied PFAS concentration observed between Indo-Pacific humpback dolphins (*S. chinensis*) and finless porpoises (*N. phocaenoides*) (Yeung et al., 2009). Genus-specific differences have been reported between five odontocete species, with *Tursiops* displaying PFAS levels three times higher than all other species (López-Berenguer et al. (2020). We argue that there is a need for classifying populations nutritional niches when determining contamination risks to compare species-specific niches that may explain the variations between PFAS concentrations as presented in these results (Stockin et al., 2023).

Across the highest ranked models (Table 1) for ∑PFAS, ∑PFCAs and PFOS, our results show coastal species demonstrated the highest predicted PFAS contamination. These included the Indo-Pacific humpback dolphin (*S. chinensis*), bottlenose dolphins (*Tursiops* spp.) and the Indo-Pacific finless porpoise (*N. phocaenoides*). Notably, the Indo-Pacific humpback dolphin and Indo-Pacific finless porpoise were the sole species representing the *Sousa* and *Neophocaena* genus groups, respectively (Table S9, Table S12). These results indicate that, at a global scale, coastal environments likely expose these species to higher contamination. Moreover, the Indo-Pacific humpback dolphin and Indo-Pacific finless porpoise are species resident to China, which has been the primary manufacturer of long-chain PFAS since stronger regulation of production of PFAS in countries such as Europe and America (Li et al., 2015; Wang et al., 2014). There could be direct links to pollution from high industrialisation and urbanisation levels in the Guangdong–Hong Kong–Macao Greater Bay Area, which *Sousa* and *Neophocaena* primarily inhabit (Wang et al., 2021; Yang et al., 2019). Here, PFAS concentration in surface water in the Yangtze River and Pearl River Estuary are among the highest worldwide (Pan et al., 2018; Wang et al., 2019). It is likely that, in addition to direct absorption of PFAS from the water column, the high concentrations in these species result from bioaccumulation and biomagnification processes, whereby prey are contaminated due to their close proximity to anthropogenic sources (Johnson-Restrepo et al., 2005; Suedel et al., 1994).

In addition to ecological and geographical factors, it is likely other biological factors influence PFAS in odontocetes. For example, Stockin et al. (2025) assessed PFAS in livers across 16 cetacean species in New Zealand and found habitat did not influence PFAS levels. We suggest physiological factors could increase levels observed in certain species, such as higher concentrations in *Tursiops*. For example, in an aquarium-based study, bottlenose dolphins (*T. truncatus*) had more than double the amount of PFAS concentration than killer whales, even though both species had the same water supply (Lemos et al., 2024). It is suggested lower PFAS levels in killer whales could be due to differences in prey supply or a potential ‘dilution effect’, where contamination decreases with the size of an animal (Kamunde & Wood, 2003; Lemos et al., 2024; Sciancalepore et al., 2021).

Comparing PFAS levels in different ecotypes and from different regions would help to better understand the influence of ecology or the impact of a potential dilution effect (Fig. 1, Table 2). Further, while metabolic rates in odontocetes remain largely unknown (Noren & Rosen, 2023), phylogenetic differences may affect the transformation ability of PFAS across species (Fig. 1). A comparison of PFOSA-to-PFOS ratios in harbour porpoises (*P. phocoena*) and white-beaked dolphins (*L. albirostris*), two sympatric species with similar diets, showed marked interspecific differences in PFAS levels and profiles, suggesting species-specific metabolism or exposure pathways (Galatius et al., 2013). Understanding of interspecific differences in metabolism and how these may influence PFAS elimination rates across species may increase our understanding of PFAS contamination in odontocetes.

### 4.2 Spatial-temporal effects on PFAS contamination

Varying PFAS contamination levels across ocean and sea basins were observed across the highest-ranked models for each response variable (∑PFAS, ∑PFCAs and PFOS) (Fig. 1). The results may in part, reflect the differences in production, use and regulatory histories of PFAS worldwide (Paul et al., 2009; Tansel, 2024). For example, we observed lower ∑PFAS and PFOS concentrations in the Mediterranean Sea comparative to other ocean regions (Fig. 1), which may be consistent with Europe’s restrictions and the cessation of PFOS production by major PFAS producing companies such as 3M (Paul et al., 2009). Comparatively, there were elevated ∑PFCAs in the Mediterranean Sea that may mirror the increased reliance on carboxylate substitutes such as PFNA, following PFOS phase-outs (Dean et al., 2020; Langenbach & Wilson, 2021). Similar results have been reported in Scopoli’s shearwaters (*Calonectris dimedea*) in the Western Mediterranean Sea, where PFCAs exceeded PFOS levels following the regulation of PFOS under the Stockholm Convention in 2009 (Lu et al., 2024; UNEP, 2020). A study on striped dolphins (*Stenella coeruleoalba*) in the Mediterranean Sea similarly reported increasing proportions of PFCAs over time alongside stabilising PFOS concentrations, reflecting reduced inputs of PFOS due to regulatory efforts (Garcia-Garin et al., 2023). Despite this stabilisation, PFOS remains one of the most prevalent and bioaccumulative PFAS compounds in odontocetes (Garcia-Garin et al., 2023), however these results highlight the importance of regulation to limit further accumulation.

For example, PFAS levels in the Pacific Ocean, displayed the highest contamination across all final models in our study (Fig. 1), which aligns with ongoing production of PFAS including, long-chain PFOS, in parts of Asia, particularly China (Wang et al., 2014).

Furthermore, the mixed results reported in previous marine mammal research (Law, 2014), likely explains the minimal increase of PFAS contamination through the years, as observed in our ∑PFAS and ∑PFCAs models (Table 2, Fig. 2), and our exclusion of year as a factor in our final PFOS model. Increasing concentrations of carboxylates were observed from the 1980s to the early 2000s in harbor porpoises (*P. phocoena*) in the Baltic Sea and Iceland (Huber et al., 2012), as well as in the Danish North Sea (Galatius et al., 2011). Similar increasing trends were observed in harbor porpoises, long-finned pilot whales (*Globicephala melas*), along with several other marine mammals, from 1984 to 2009, in the north Atlantic and west Greenland (Rotander et al., 2012). However, mixed trends were observed in PFOS, including levels of sulfonates declining in harbor porpoises in the Baltic and North Seas between 1991 to 2008 (Huber et al., 2012). In Rotander et al. (2012), sampling from 1984 to 2009 showed that PFOS remained stable and generally displayed a downward trend in marine mammals globally, although PFCAs continued to increase (Houde et al., 2011). In contrast, a considerable 10-fold increase in PFOS contamination was observed in melon headed whales (*Peponocephala electra*) in Japan from 1982 to 2002 (Hart et al., 2008).

**Figure 2.**
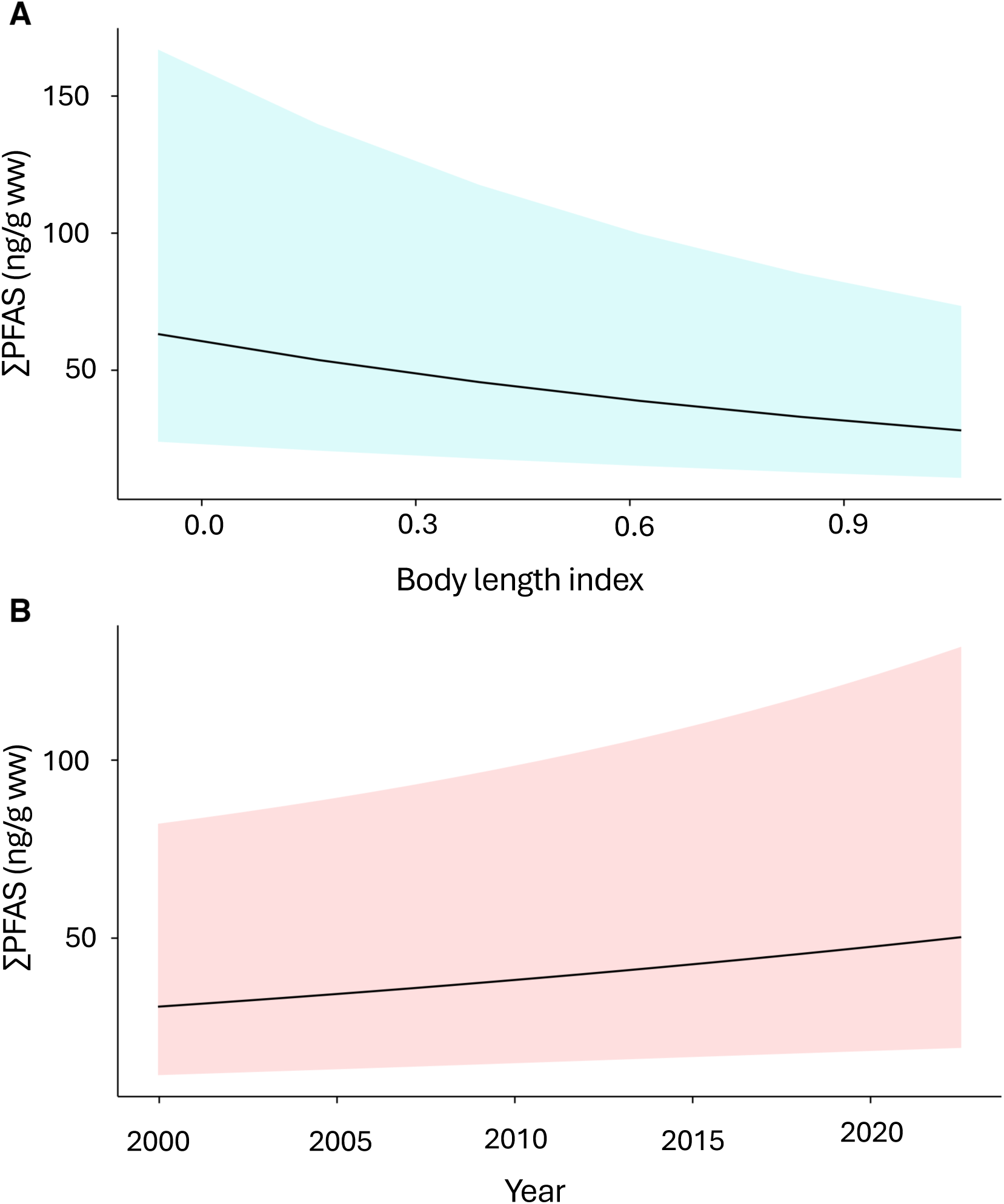
Predicted ∑PFAS concentrations (ng/g wet weight) from generalised linear mixed models (GLMMs), shown in relation to (A) body length index (proxy for life stage) displaying ∼15% decrease in contamination with increasing life stage and, (B) sampling year for odontocetes, with contamination increasing ∼ 14% over time (n = 713). Shaded areas represent 95% confidence.

These findings show PFAS compounds are ubiquitous in the environment and are likely extremely persistent due to their resistance to breakdown (Brunn et al., 2023). The observed increase in PFAS contamination may be linked to delayed transport processes such as the movement of PFAS from soils to water systems or atmospheric deposition, which can have a lag time of up to 10 years (Lam et al., 2016; Law, 2014). These factors may help explain the elevated ∑PFAS, PFOS and ∑PFCAs observed in other regions such as Oceania (Fig. 1) which remain unexplained despite similar regulatory frameworks to Europe and America, and no known large-scale manufacturing (Brennan et al., 2021). The predictor variables in our ∑PFAS model explained 76% of the deviance of PFAS contamination (Table S15), therefore considerable variation remains unexplained. It would be useful to link PFAS concentrations to key point and non-point sources, and consider abiotic transport mechanisms such as upwelling, downwelling, currents, and weather patterns in future, to help provide a more complete understanding of PFAS distribution and dynamics. Furthermore, future efforts should address major geographical gaps, such as underrepresented regions in Antarctica and Africa, and in highly urbanised and industrialised regions such as India and Indonesia.

Additionally, improvements in data transparency and consistency in data handling across the literature on PFAS contamination is necessary. Of the 42 papers initially identified, only 18 could be included in this analysis, reducing the usable sample size from over 1,300 to 713.

This resulted in the loss of data from different regions and reduced the explanatory power of global PFAS concentration models. To minimize such losses in future work, improved data management practices, standardized reporting, and greater accessibility of datasets and metadata are strongly recommended to facilitate large-scale comparative analyses.

### 4.3 Varying effects across PFAS compounds

Differences in spatial results across our three models highlights the importance of quantifying a broad suite of PFAS compounds. In general, PFOS has larger absolute concentrations and therefore, disproportionately influences ∑PFAS estimates, partly explaining why regions with high ∑PFAS corresponded to areas with elevated PFOS (Fig. 1, S16, S17). While our study assesses more compounds than current published global analyses on PFAS contamination in wildlife (Armitage et al., 2009), providing the first quantitative global analysis on odontocetes, it is recommended that future studies expand the range of compounds assessed, in order to gain more knowledge about the distribution, prevalence, and effects of many other PFAS compounds. Across studies, there was considerable variation in the types of PFAS compounds assessed, with at least 57 individual PFAS compounds or groups identified in odontocetes. However, only five of these compounds were consistently selected for analysis across all studies and thus, included in our analysis. Furthermore, our results indicate that PFCA and PFOS levels show different relationships with predictor variables; for example, sex had no effect on carboxylate levels, while time had no effect on sulfonates (PFOS) levels in odontocetes globally (Table 1).

### 4.4 Influence of life stage on PFAS contamination

We found life stage, as estimated by a standardized index of total body length respective to each species, to predict the amount of ∑PFAS concentration in odontocetes, with ∑PFAS concentrations decreasing with stage of life (Fig. 2, Table 1). These findings are in agreement with higher concentrations of PFAS reported in juveniles compared to mature individuals (Fair et al., 2012; Foord et al., 2024; Galatius et al., 2011; Stockin et al., 2021). Younger individuals often exhibit higher PFAS contamination from transferal across the placental barrier and through suckling (Fair et al., 2012; Galatius et al., 2011; Lynch et al., 2019; Stockin et al., 2021; Stockin et al., 2025). There is increasing evidence of offloading from mothers to their offspring including in tucuxi dolphins (*Sotalia guianensis*) from Brazil (Dorneles et al., 2008) in beluga whales (*Delphinapterus leucas*) from the Arctic (Kelly et al., 2009), and in captive common bottlenose dolphins (*T. truncatus*) in America (Lemos et al., 2024). PFAS contaminants have been detected in the milk of common bottlenose dolphins in America (Houde et al., 2006) and in Australia (Gaylard, 2016), where a stomach content sample from a *Tursiops* calf days old was analysed, which presented high levels with 890 ng/g of PFOS. Additionally long-term assessments of mother-calf pairs of killer whales and common bottlenose dolphins revealed high PFAS concentrations in newborn calves, providing evidence of PFAS transfer in utero, with further increased levels during the suckling period (Lemos et al., 2024). The results presented here did not include fetuses, and 14 samples were excluded from calves due to unidentified sex. Accordingly, our results may underestimate the contaminant levels in calves. We recommend investigation into the specific processes causing in-utero transfer of PFAS, including differences between PFAS compounds for future research and the potential impacts to development of odontocetes from this early life exposure of contaminants (Galatius et al., 2011; Lemos et al., 2024; Stockin et al., 2021).

However, offloading of contaminants from mother to calf does not explain reduction of PFAS contamination for any male individuals, which challenges the hypothesis of bioaccumulation, where contamination is expected to increase throughout an animal’s life (Bryan & Darracott, 1979; Mackay & Fraser, 2000). Instead, our results could support the idea of a potential dilution effect, where contaminants may dilute with body size or growth. While this process is yet to be tested for PFAS accumulation, it has been observed for copper exposure in rainbow trout (*Oncorhynchus mykiss*) (Kamunde & Wood, 2003) and could be important to consider in future research. As suggested by Gui et al. (2019), increased elimination of PFAS in adults, could be related to metabolic transform abilities, as it is likely calves do not have developed metabolic mechanisms (Sly & Flack, 2008; Xie et al., 2022; Zhang et al., 2021). However, it contradicts findings that suggest metabolic rate decreases within an odontocetes life (as reviewed in Noren & Rosen, 2023).

### 4.5 Differences in PFAS burden between sexes

As males reported higher PFAS concentrations compared to females (Table 2), sex may play a role in PFAS biotransformation or offloading capacity in odontocetes. These findings could be attributed to the offloading of contaminants from mature females to their calves, as suggested by Stockin et al. (2021). As a result, sexually mature females that have recently given birth may have decreased contamination levels due to offloading contaminants to their young, while conversely, newborn delphinids may exhibit increased contamination, as found in this study (Fig. 2) (Fair et al., 2012; Lynch et al., 2019). In Lemos et al. (2024), higher PFAS concentrations where reported in male bottlenose dolphins compared to female counterparts.

However, in killer whales, this difference was evident only when comparing mature males with non-lactating mature females, after excluding immature, pregnant, and lactating individuals, suggesting that maternal offloading to calves substantially influence PFAS burdens in odontocetes. This may partly explain why we detected no sex-based differences in our ∑PFCAs model, as we were unable to account for differences among immature, pregnant, and lactating individuals, which could contribute to variations in PFAS concentrations.

The higher ∑PFAS contamination observed in males, compared to females, may be due to differences in the ability to eliminate contaminants between sexes in odontocetes. For instance, sex hormones play an important role in biological elimination of contaminants through urine (Han et al., 2012). A study on rats (*Rattus rattus*) compared renal clearance of PFOA with two hormone treated males, including 17β-estradiol and testosterone, resulting in 17β-estradiol males increasing their elimination of PFOA, matching that of females, and high testosterone males with reduced removal of contaminants (Han et al., 2012; Vanden Heuvel et al., 1992). Future studies should further investigate the relationship between PFAS contamination and sex hormones, including across different PFAS compounds, to better understand the mechanisms underlying potential sex-specific bioelimination processes.

## Conclusion

PFAS contamination in odontocetes is influenced by species, location, age, sex, and time. Among these, genus was the most influential factor, highlighting the importance of interspecific variation in understanding large-scale contamination patterns. Genus and geographic differences likely reflect variation in pollution exposure across countries and ecological niches, emphasising the role of ecological parameters such as diet, physiology, and trophic interactions. The significant effect of life stage, where PFAS concentrations decreased with age, is consistent with maternal offloading to calves, which may also suggest partial elimination of contaminants over an individual’s lifetime. Lower PFAS levels in females compared to males likely result from reproductive offloading and sex-specific hormonal influences on elimination rates. The temporal increase in contamination levels highlights the pervasive and persistent nature of PFAS across marine ecosystems.

Collectively, these findings emphasise the need for future research to identify the physiological and ecological mechanisms driving PFAS accumulation and elimination globally, including the roles of maternal transfer, metabolic processes, and endocrine regulation.

## CRediT authorship contribution statement

**Lavinia Stokes**: Data curation, Methodology, Formal analysis, Investigation, Visualization, Writing – original draft.

**Katharina J. Peters:** Conceptualization, Methodology, Formal analysis, Visualization, Resources, Supervision, Project administration, Writing – review and editing.

**Karen A. Stockin**: Conceptualization, Resources, Funding acquisition, Supervision, Writing – review and editing.

**Frédérik Saltré:** Formal analysis, Methodology, Visualization, Writing – review and editing.

**Gavin Stevenson:** Formal analysis, Writing – reviewing and editing. **Jesuina De Araujo:** Formal analysis, Writing – review and editing.

## Acknowledgement

We thank Victor Peddemors and his team at the Department of Primary Industries in Sydney for providing samples. We are grateful to all the researchers who provided their raw data and associated information.

## Data availability statement

Data and code that support our findings are openly available at github.com/Lavinia-Stokes/Global-PFAS-Odontocetes

