## Supplementary document for "Global patterns and predictors of PFAS contamination in odontocetes"

***S1**. Data collection of liver samples from deceased dolphins in Australian waters.*

Liver samples were collected opportunistically from two Indo-Pacific bottlenose dolphin (*Tursiops aduncus*) and two common dolphin (*Delphinus delphis*) that had died following incidental entanglement in shark nets in southeastern NSW between 2017 and 2019 (see S2 and S4). Total body length (TBL) and sex was assessed by Vic Peddemors during necropsy (S4) however, individual NAR190126 could not be sexed as the specimen was a calf and lacked fully developed sexual organs. Liver samples were extracted from each animal, wrapped in aluminium foil, placed in zip lock plastic bags and stored at (-20°C).

We subsampled the liver specimens by extracting a section from the central section of the middle part of the sample using a stainless-steel knife on a glass chopping board. Samples were weighed to 30 g using a glass cooking scale and placed in high density polyethylene sample containers. Subsamples were kept at -20°C until transferred to the analytical testing service, National Measurement Institute in North Ryde, Sydney.

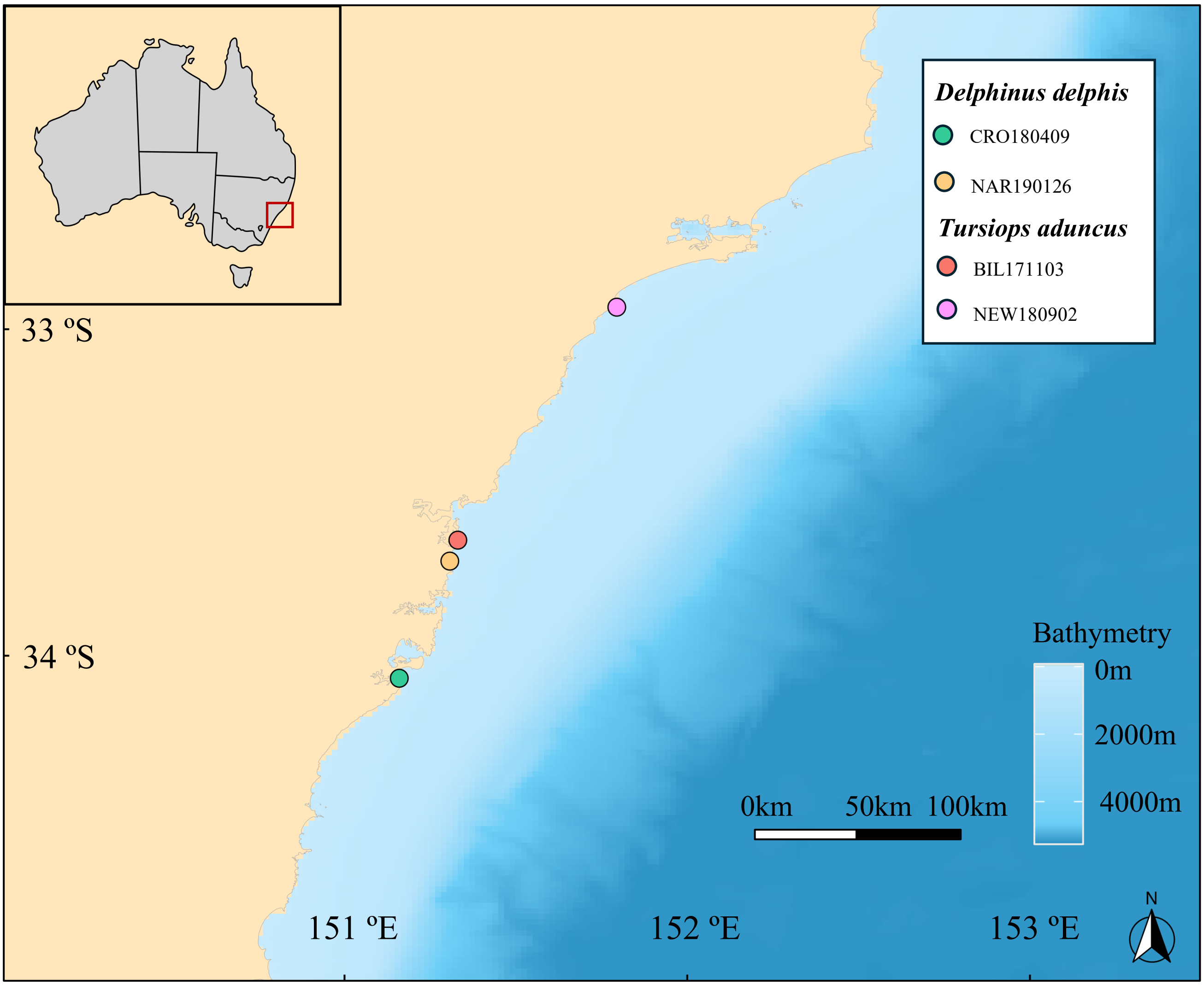


**S2**. Entanglement locations for four dolphins along the coast of NSW, Australia from 2017 to 2019. Each individual/sample is represented by a different coloured circle. Bathymetry (Deane, 2019), is illustrated by the blue gradient scale, where darker shades represent deeper waters and lighter shades represent shallower waters.

***S3**. Analytical methodology*

Dolphin liver tissue samples were analysed for PFAS by National Measurement Institute (NMI), North Ryde, Sydney, NSW, Australia. To quantify PFAS contamination in dolphin liver samples, NMI used High-Performance Liquid Chromatography coupled with Triple Quadrupole Mass Spectrometry (LCMS/MS, ). Isotopically labelled extraction internal standards (Wellington Laboratories, Canada) were added to the samples to correct for any losses during sample preparation and to serve as references for quantifying PFAS concentrations. Alkaline methanol (0.01N Potassium Hydroxide in Methanol, ChemSupply AR grade, and Merck LCMS grade respectively) was used to extract the PFAS from the homogenized biota using overnight tumbling, then the extract was separated by centrifuging and purified with activated carbon (Envi-CARB 120-400 Mesh SPE, 1g, Supelco). Further isotopically labelled standards were added to act as injection standards. The extracted sample was then subjected to liquid chromatography (SCIEX Exion with C18, Aquity BEH XBridge, 2.1 x 100 mm x 1.7 µm, 130 Å, Waters, USA) to separate the PFAS compounds based on their chemical properties and retention times. Following this, the compounds were analysed using electrospray mass spectrometry (SCIEX 6500+, USA), where PFAS molecules were broken into ionised fragments and their mass-to-charge ratios were measured. Multiple reaction monitoring was used to analyse two characteristic ions for each PFAS compound to enhance specificity. Analytes were identified by detecting specific ions within the selected ion monitoring windows, and quantification was achieved by comparing the detected amounts to the labelled internal standards, PFHxS and PFOS calibration standards contained branched and linear isomers, which were calculated separately and summed for reporting. All results were reported on a wet weight basis (w/w) in mg/kg. A total of 33 PFAS compounds were tested, including 13 perflouroalkane carboxylic acids, seven perfluoroalkane sulfonic acids, seven perfluoroalkyl sulfonamides, four fluorotelomer sulfonic acids, a phosphate ester and an unsaturated telomer acid, refer to S4.

**S4.** Five PFAS compounds consistently reported in global studies on PFAS contamination in odontocetes, including the chemical names and subgroups.

| **Abbreviation** | **Chemical Name** | **Subgroup** |
|---|---|---|
| *PFNA* | Perfluorononanoic acid | Carboxylate |
| *PFDA* | Perfluorodecanoic acid | Carboxylate |
| *PFUnDA* | Perfluoroundecanoic acid | Carboxylate |
| *PFDoDA* | Perfluorododecanoic acid | Carboxylate |
| *PFOS* | Perfluorooctanesulfonic acid | Sulfonate |

**S5.** Identification information and summary of PFAS concentration in four dolphin samples from southeastern NSW, Australia. Values above limit of reporting (LOR) are in ng/g w/w.

| **Dolphin ID** | ***NAR190126*** | ***NEW180902*** | ***BIL171103*** | ***CRO180409*** | |
|---|---|---|---|---|---|
| **Species** | *Delphinus delphis* | *Tursiops aduncus* | *T. aduncus* | *D. delphis* | |
| **Sex** | N/A | M | F | F | |
| **Total body length** | 1000 mm | 2211 mm | 1691mm | 2402 mm | |
| **Location** | Narrabeen, NSW | Newcastle, NSW | Bilgola, NSW | Cronulla, NSW | |
| **Date** | 26/1/2019 | 2/9/2018 | 3/11/2017 | 9/4/2018 | |
| | | | | | **LOR** |
| PFBA (375-22-4) | <0.001 | <0.001 | <0.001 | <0.001 | 0.001 |
| PFPeA (2706-90-3) | <0.0005 | <0.0005 | <0.0005 | <0.0005 | 0.0005 |
| PFHxA (307-24-4) | <0.0005 | <0.0005 | <0.0005 | <0.0005 | 0.0005 |
| PFHpA | 1.8 | 0.76 | 1 | 0 | 0.0005 |
| PFOA | 4 | 0.63 | 0.67 | 0 | 0.0003 |
| PFNA | 13 | 4.9 | 11 | 1.5 | 0.0005 |
| PFDA | 20 | 12 | 19 | 5.1 | 0.0005 |
| PFUnDA | 15 | 5.1 | 15 | 5.6 | 0.0005 |
| PFDoDA | 11 | 6.6 | 13 | 5.5 | 0.0005 |
| PFTrDA | 0.51 | 1.4 | 2.5 | 0.6 | 0.0005 |
| PFTeDA | 1.2 | 3.7 | 4.7 | 1.8 | 0.001 |
| PFBS | 0 | 0 | 0 | 0 | 0.0005 |
| PFPeS | 0 | 0 | 0 | 0 | 0.0005 |
| PFHxS | 5.3 | 5.4 | 2.5 | 0 | 0.0005 |
| PFHpS | 1.7 | 2 | 0.6 | 0 | 0.0005 |
| PFOS | 210 | 410 | 150 | 35 | 0.0003 |
| PFNS | 0.34 | 0.67 | 0.38 | 0 | 0.0003 |
| PFDS | 1.7 | 1.9 | 1.6 | 0.65 | 0.0003 |
| PFOSA | 9.6 | 13 | 4.1 | 7.2 | 0.001 |
| N-MeFOSA | 0 | 0 | 0 | 0 | 0.0005 |
| N-MeFOSAA | 0.58 | 0.5 | 0.88 | 0 | 0.0005 |
| N-EtFOSAA | <0.0005 | <0.0005 | <0.0005 | <0.0005 | 0.0005 |
| N-MeFOSE | <0.005 | <0.005 | <0.005 | <0.005 | 0.005 |
| 4:2 FTS | <0.0005 | <0.0005 | <0.0005 | <0.0005 | 0.0005 |
| 6:2 FTS | <0.0005 | <0.0005 | <0.0005 | <0.0005 | 0.0005 |
| 8:2 FTS | <0.0005 | <0.0005 | <0.0005 | <0.0005 | 0.0005 |
| 10:2 FTS | <0.0005 | <0.0005 | <0.0005 | <0.0005 | 0.0005 |
| HFPO-DA | <0.0005 | <0.0005 | <0.0005 | <0.0005 | 0.0005 |
| 9Cl-PF3ONS | <0.0005 | <0.0005 | <0.0005 | <0.0005 | 0.0005 |
| 11Cl-PF3OUdS | <0.0005 | <0.0005 | <0.0005 | <0.0005 | 0.0005 |
| ADONA | <0.005 | <0.005 | <0.005 | <0.005 | 0.005 |

| | | | | | |
|---|---|---|---|---|---|
| PFDoS | <0.001 | <0.001 | <0.001 | <0.001 | 0.001 |
| PFPrS | <0.001 | <0.001 | <0.001 | <0.001 | 0.001 |
| **∑PFCAs** | 66.51 | 35.09 | 66.87 | 20.1 | |
| **∑PFSAs** | 219.04 | 419.97 | 155.08 | 35.65 | |
| **∑Precursors** | 10.18 | 13.5 | 4.98 | 7.2 | |
| **∑PFAS** | 295.73 | 468.56 | 226.93 | 62.95 | |

**S6.** *Composition and profiles of PFAS burden in samples from Australia*

A total of 15 PFAS compounds were detected from the 33 compounds analysed. Of these, 7 long- chain compounds (chemicals with a carbon chain >8), were found in all samples, including carboxylic acids, (PFNA, PFDA, PFUnDA, PFDoDA, PFTeDA), sulfonic acids (PFOS, PFDS) as well as one precursor (PFOSA). Other carboxylic acids (PFHpA, PFOA) and sulfonic acids, (PFHxS, PFHpS, PFNS) were detected in all samples except for D. *delphis* sample CRO180409 which had the overall lowest ∑PFAS. Other combinations included the compound PFTrDA, that was found only in CRO180409 and BIL171103, while the precursor compound N.MeFOSAA was only present in NAR190126 and BIL171103.

Overall, PFSAs exhibited the highest burden compared to other PFAS classes across all four samples, primarily due to PFOS, which presented the highest composition across all samples. Sample NEW180902, an Indo-Pacific bottlenose dolphin (T. *aduncus*) from Newcastle and the only known male amongst the four samples, noticeably displayed the highest ∑PFAS burden (∑PFCAs = 35.09, ∑PFSAs = 419.97, ∑Precursors = 13.5). Sample CRO180409, a female common dolphin (D. *delphis*) which was the largest of all four animals, had the lowest ∑PFAS burden (∑PFCAs = 20.1, ∑PFSAs = 35.65, ∑Precursors = 7.2).

Across the two samples for each species, the Indo-Pacific bottlenose dolphin had higher ∑PFAS concentration than the common dolphin (mean ± SD = 347.75 ± 170.86 ng/g ww and 179.34 ± 164.6 ng/g ww, respectively).

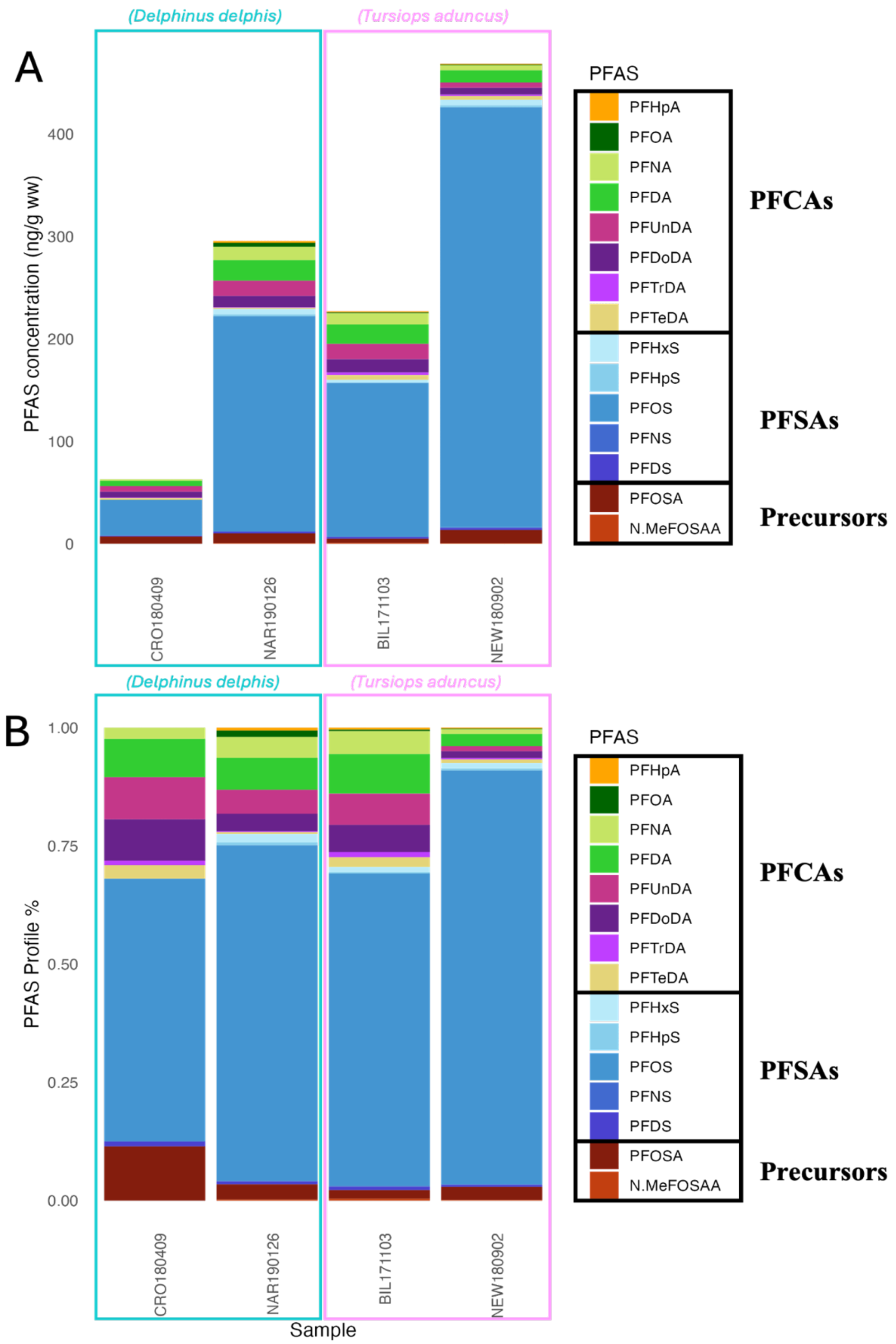


**S7.** Hepatic PFAS concentrations common dolphins (*D. delphis*; n = 2, blue boxes) and Indo-Pacific bottlenose dolphins (*T. aduncus*; n = 2, pink boxes), entangled in shark nets in NSW between 2017 and 2019. Panel A shows the total PFAS concentration for each sample.

**S9.** Minimum and maximum known total body lengths of odontocetes assessed in this study. All lengths are in metres.

| **Species** | **Minimum calf length** | **Maximum length** | | **Reference** |
|---|---|---|---|---|
| | | **female** | **male** | |
| Arnoux's beaked whale (*B. arnuxii*) | 4 | 9.75 | 9.3 | (Ellis & Mead, 2017); National Strandings Database, Department of Conservation, New Zealand; CERG-Massey University Biology Database |
| Hector's dolphin (*C. hectori*) | 0.5 | 1.53 | 1.4 | (Casano-Bally, 2023; Stockin et al., 2025; Webster et al., 2010); National Strandings Database, Department of Conservation, New Zealand; CERG-Massey University Biology Database |
| beluga whale (*D. leucas*) | 1.3 | 4.3 | 5.5 | (Würsig et al., 2017) |
| common dolphin (*D. delphis*) | 0.76 | 2.6 | 2.4 | (Palmer et al., 2023; Perrin, 2018; Ross, 2006) |
| long-finned pilot whale (*G. melas*) | 1.38 | 6 | 7.2 | (Ross, 2006; Würsig et al., 2017) |
| Risso's dolphin (*G. griseus*) | 1.1 | 4.3 | 3.7 | (Würsig et al., 2017); Whales and Dolphins of the Marianas, NOAA) |
| pygmy sperm whale (*K. breviceps*) | 1.2 | 3.8 | 3.4 | (Würsig et al., 2017); Department of Conservation, New Zealand; CERG-Massey University Biology Database |
| white-beaked dolphin (*L. albirostris*) | 0.95 | 2.7 | 3.1 | (Kinze, 2009; Lauria et al., 2024; Würsig et al., 2017) |
| hourglass dolphin (*L. cruciger*) | 0.9 | 1.83 | 1.87 | (Goodall et al., 1997); National Strandings Database, Department of Conservation, New Zealand; CERG-Massey University Biology Database |
| dusky dolphin (*L. obscurus*) | 0.9 | 1.95 | 1.92 | (Cipriano, 1992; Poelijoe, 2024); National Strandings Database, Department of Conservation, New Zealand; CERG-Massey University Biology Database |
| Gray's beaked whale (*M. grayi*) | 2.1 | 5.9 | 6.2 | (Thompson et al., 2013); Department of Conservation, New Zealand; CERG-Massey University Biology Database |

| | | | | |
|---|---|---|---|---|
| strap-toothed beaked whale (*M. layardii*) | 2.5 | 6.2 | 5.9 | Department of Conservation, New Zealand; CERG-Massey University Biology Database |
| Indo-Pacific finless porpoise (*N. phocaenoides*) | 0.6 | 1.76 | 1.9 | (Wang et al., 2021; Würsig et al., 2017) |
| killer whale (*O. orca*) | 2.02 | 9.75 | 9.8 | (Ross, 2006; Stockin et al., 2025) |
| melon-headed whale (*P. electra*) | 1 | 2.75 | 2.7 | (Hart et al., 2008; Würsig et al., 2017) |
| spectacled porpoise (*P. dioptrica*) | 0.46 | 2.04 | 2.24 | (Dellabianca et al., 2025); Department of Conservation, New Zealand; CERG-Massey University Biology Database |
| sperm whale (*P. macrocephalus*) | 3.5 | 12 | 18 | (Giorli & Goetz, 2020); Department of Conservation, New Zealand; CERG-Massey University Biology Database |
| Franciscana dolphin (*P. blainvillei*) | 0.7 | 1.77 | 1.74 | (Crespo, 2018; Würsig et al., 2017) |
| false killer whale (*P. crassidens*) | 1.2 | 5.06 | 5.96 | (Ross, 2006) |
| Guiana dolphin (*S. guianensis*) | 0.9 | 2.2 | 2.2 | (Würsig et al., 2017) |
| Indo-Pacific humpback dolphin (*S. chinensis*) | 0.9 | 2.7 | 2.8 | (Lam et al., 2016; Würsig et al., 2017) |
| striped dolphin (*S. coeruleoalba*) | 0.8 | 2.5 | 2.6 | (Ross, 2006; Würsig et al., 2017) |
| Indo-Pacific bottlenose dolphin (*T. aduncus*) | 0.83 | 2.7 | 2.7 | (Ross, 2006; Würsig et al., 2017) |

| | | | | |
|---|---|---|---|---|
| Burrunan dolphin (*T. australis*) | 0.83 | 2.78 | 2.78 | (Charlton-Robb et al., 2011; Ross, 2006; Würsig et al., 2017) |
| common bottlenose dolphin (*T. truncatus*) | 0.9 | 3.6 | 3.9 | (Crowe et al., 2025; Würsig et al., 2017); National Strandings Database, Department of Conservation, New Zealand; CERG-Massey University Biology Database |
| Cuvier's beaked whale (*Z. cavirostris*) | 2.5 | 6.2 | 6.2 | (MacLeod, 2005; Nishiwaki & Oguro, 1972); National Strandings Database, Department of Conservation, New Zealand; CERG-Massey University Biology Database |
| pygmy killer whale (*F. attenuata*) | 0.9 | 2.6 | 2.5 | (Boys et al., 2023); Department of Conservation, New Zealand; CERG-Massey University Biology Database |
| Longman's beaked whale (*I. pacificus*) | 2.91 | 6.48 | 6.08 | (Ross, 1984; Wang et al., 2006) |
| dwarf sperm whale (*K. sima*) | 1 | 2.86 | 2.62 | (Ross, 2006; Würsig et al., 2017) |
| Blainville's beaked whale (*M. densirostris*) | 1.9 | 4.71 | 5.8 | (Cooke & Klinowska, 1991; Würsig et al., 2017) |
| pantropical spotted dolphin (*S. attenuata*) | 0.8 | 3.4 | 2.57 | (Ross, 2006; Würsig et al., 2017) |
| spinner dolphin (*S. longirostris*) | 0.7 | 2.07 | 2.35 | (Ross, 2006; Würsig et al., 2017) |
| rough-toothed dolphin (*S. bredanensis*) | 1 | 2.7 | 2.83 | (Würsig et al., 2017) |

**S10.** Model Justification

To determine the appropriate model structure for ∑PFAS contamination, we calculated the intraclass correlation coefficient and the design effect for the grouping factors, genus and location. For genus, an intraclass correlation coefficient of 0.069, indicated that ~6.9% of the total variation in ∑PFAS concentration was due to differences between genera, while the remaining 93.1% reflected individual variation within clusters. The design effect of 3.269 suggests the grouping structure substantially inflates standard errors by more than 3-fold compared to a simple random sample. The intraclass correlation coefficient was lower for location (i.e., 0.035), explaining ~3.5% of the total variation, whereas large group sizes increased the design effect (i.e., ~5.143). Using the same approach, results showed that for both ∑PFCAs and PFOS, intraclass correlation coefficients varied between 1.3% and 9.5%, while the design effect ranged from 1.69 to 6.47. These results indicate that, despite modest between-group variance, clustering by genus and location inflated standard errors, justifying their inclusion as random effects to ensure valid inference and robust fixed-effect estimates.

**S11.** List of species groupings used in generalised linear model to analyse global PFAS concentrations in odontocetes. Species were grouped by taxonomic level, resulting in 21 groups spanning 33 odontocete species.

| Group/Genus | Species | Common Name |
|---|---|---|
| Beaked whales | *Berardius arnuxii* | Arnoux's beaked whale |
| | *Indopacetus pacificus* | Longman's beaked whale |
| | *Mesoplodon densirostris* | Blainville's beaked whale |
| | *M. grayi* | Gray's beaked whale |
| | *M. layardii* | strap-toothed beaked whale |
| | *Ziphius cavirostris* | Cuvier's beaked whale |
| *Cephalorhynchus* | *Cephalorhynchus hectori* | Hector's dolphin |
| *Delphinapterus* | *Delphinapterus leucas* | beluga whale |
| *Delphinus* | *Delphinus delphis* | common dolphin |
| *Feresa* | *Feresa attenuata* | pygmy killer whale |
| *Globicephala* | *Globicephala melas* | long-finned pilot whale |
| *Grampus* | *Grampus griseus* | Risso's dolphin |
| *Kogia* | *Kogia breviceps* | Pygmy sperm whale |
| | *K. sima* | dwarf sperm whale |
| *Lagenorhynchus* | *Lagenorhynchus albirostris* | white-beaked dolphin |
| | *L. cruciger** | hourglass dolphin |
| | *L. obscurus** | dusky dolphin |
| *Neophocaena* | *Neophocaena phocaenoides* | Indo-pacific finless porpoise |
| *Orcinus* | *Orcinus orca* | killer whale |
| *Peponocephala* | *Peponocephala electra* | melon-headed whale |
| *Phocoena* | *Phocoena dioptrica* | spectacled porpoise |
| *Physteter* | *Physteter macrocephalus* | sperm whale |
| *Pontoporia* | *Pontoporia blainvillei* | Franciscana dolphin |
| *Pseudorca* | *Pseudorca crassidens* | false killer whale |
| *Sotalia* | *Sotalia guianensis* | Guiana dolphin |
| *Sousa* | *Sousa chinensis* | Indo-Pacific humpback dolphin |
| *Stenella* | *Stenella attenuata* | pantropical spotted dolphin |
| | *S. coeruleoalba* | striped dolphin |
| | *S. longirostris* | spinner dolphin |
| *Steno* | *Steno bredanensis* | rough-toothed dolphin |
| *Tursiops* | *Tursiops aduncas* | Indo-Pacific bottlenose dolphin |
| | *T. australis* | Burrunan dolphin |
| | *T. truncatus* | common bottlenose dolphin |

*since the analyses were completed, the genus of *Lagenorhynchus cruciger* and *Lagenorhynchus obscurus* have been revised (Galatius et al., 2025; Voller et al., 2019), however, for our analysis we followed the original taxonomy.

**S12.** Summary of generalised linear mixed model (GLMM) results predicting log-transformed ∑PFCAs concentrations in odontocetes (n = 713). Estimates, standard errors (SE), test statistics, and p-values are shown for fixed effects: Age Index and Year. Variance (σ) and standard deviation (SD) are reported for random intercepts associated with Ocean location and Genus group.

| Term | Estimate | SE | Statistic | p-value |
|---|---|---|---|---|
| **(Intercept)** | 2.56 | 0.49 | 5.24 | <0.0001 |
| **Age Index** | -0.19 | 0.037 | -5.13 | <0.0001 |
| **Year** | 0.38 | 0.059 | 6.42 | <0.0001 |

**S13.** Summary of generalised linear mixed model (GLMM) results predicting log-transformed PFOS concentrations in odontocetes (n = 713). Estimates, standard errors (SE), test statistics, and p-values are shown for fixed effects: Sex and Age Index. Variance (σ) and standard deviation (SD) are reported for random intercepts associated with Ocean location and Genus group.

| Term | Estimate | SE | Statistic | p-value |
|---|---|---|---|---|
| **(Intercept)** | 3.036 | 0.55 | 5.53 | <0.0001 |
| **Sex** | 0.23 | 0.11 | 2.12 | <0.01 |
| **Age Index** | -0.14 | 0.045 | -3.076 | <0.001 |

**S14.** Full model comparison results from generalised linear mixed models (GLMMs) predicting log-transformed ∑PFAS concentrations in odontocetes (n = 713), ranked by $AIC_c$. Shown are the model structure, maximum log-likelihood (LL), number of parameters (k), corrected Akaike's information criterion ($AIC_c$), change in AICc ($\Delta AIC_c$), AICc weight ($AIC_cWt$), cumulative weight (Cum.Wt), and percentage deviance explained (%DE). Fixed effects include sex (S), stage of life index (Li), and year (T); random effects include Ocean (O) and genus (G).

| Model | LL | k | $AIC_c$ | $\Delta AIC_c$ | AICcWt | Cum.Wt | %DE |
|---|---|---|---|---|---|---|---|
| **∑PFAS (n = 713)** | | | | | | | |
| S + Ai + T + (O) + (G) | -4282.55 | 8 | 8581.30 | 0.00 | 0.00 | 0.58 | 76.1% |
| S + Ai + (O) + (G) | -4284.58 | 7 | 8583.32 | 2.03 | 0.21 | 0.79 | 75.6% |
| S + Ai + T + (S*Ai) + (O) | -4282.54 | 9 | 8583.35 | 2.05 | 0.21 | 1.00 | 76.1% |
| S + (O) + (G) | -4292.76 | 6 | 8597.63 | 16.33 | 0.00 | 1.00 | 74.5% |
| (O) + (G) | -4295.08 | 5 | 8600.24 | 18.94 | 0.00 | 1.00 | 73.8% |
| Base Model | -4309.85 | 4 | 8627.75 | 46.45 | 0.00 | 1.00 | 0.0 |
| **∑PFCAs (n = 713)** | | | | | | | |
| Ai + T + (O) + (G) | -2888.77 | 7 | 5791.70 | 0.00 | 1.00 | 1.00 | 74.6% |
| Ai + (O) + (G) | -2908.48 | 6 | 5829.09 | 37.39 | 0.00 | 1.00 | 72.8% |
| (O) + (G) | -2927.72 | 5 | 5865.52 | 73.82 | 0.00 | 1.00 | 70.4% |
| S + (O) + (G) | 2927.36 | 6 | 5866.84 | 75.14 | 0.00 | 1.00 | 70.4% |

| | | | | | | | |
|---|---|---|---|---|---|---|---|
| Base PFCAs model | -2927.72 | 4 | 5923.49 | 131.78 | 0.00 | 1.00 | 0.0 |
| **PFOS (n = 713)** | | | | | | | |
| S + Ai + (O) + (G) | -3969.84 | 7 | 7953.84 | 0.00 | 0.44 | 0.44 | 81.4% |
| S + Ai + T + (O) + (G) | -3969.11 | 8 | 7954.43 | 0.59 | 0.33 | 0.77 | 81.5% |
| S + Ai + (S*Ai) + (O) + (G) | -3969.53 | 8 | 7955.27 | 1.43 | 0.22 | 0.99 | 81.4% |
| S + (O) + (G) | -3975.02 | 6 | 7962.16 | 8.32 | 0.01 | 1.00 | 80.7% |
| (O) + (G) | -3976.76 | 5 | 7963.60 | 9.76 | 0.00 | 1.00 | 80.2% |
| Base PFOS model | -4000.88 | 4 | 8009.81 | 55.96 | 0.00 | 1.00 | 0.0 |

**S15.** Variation in random intercept effects for genus groups A (black) and ocean locations B (blue) from the model predicting log ∑PFCAs concentrations. Values represent deviations from the overall intercept on the log scale, with positive values indicating higher contamination and negative values indicating lower contamination relative to the model's grand mean.

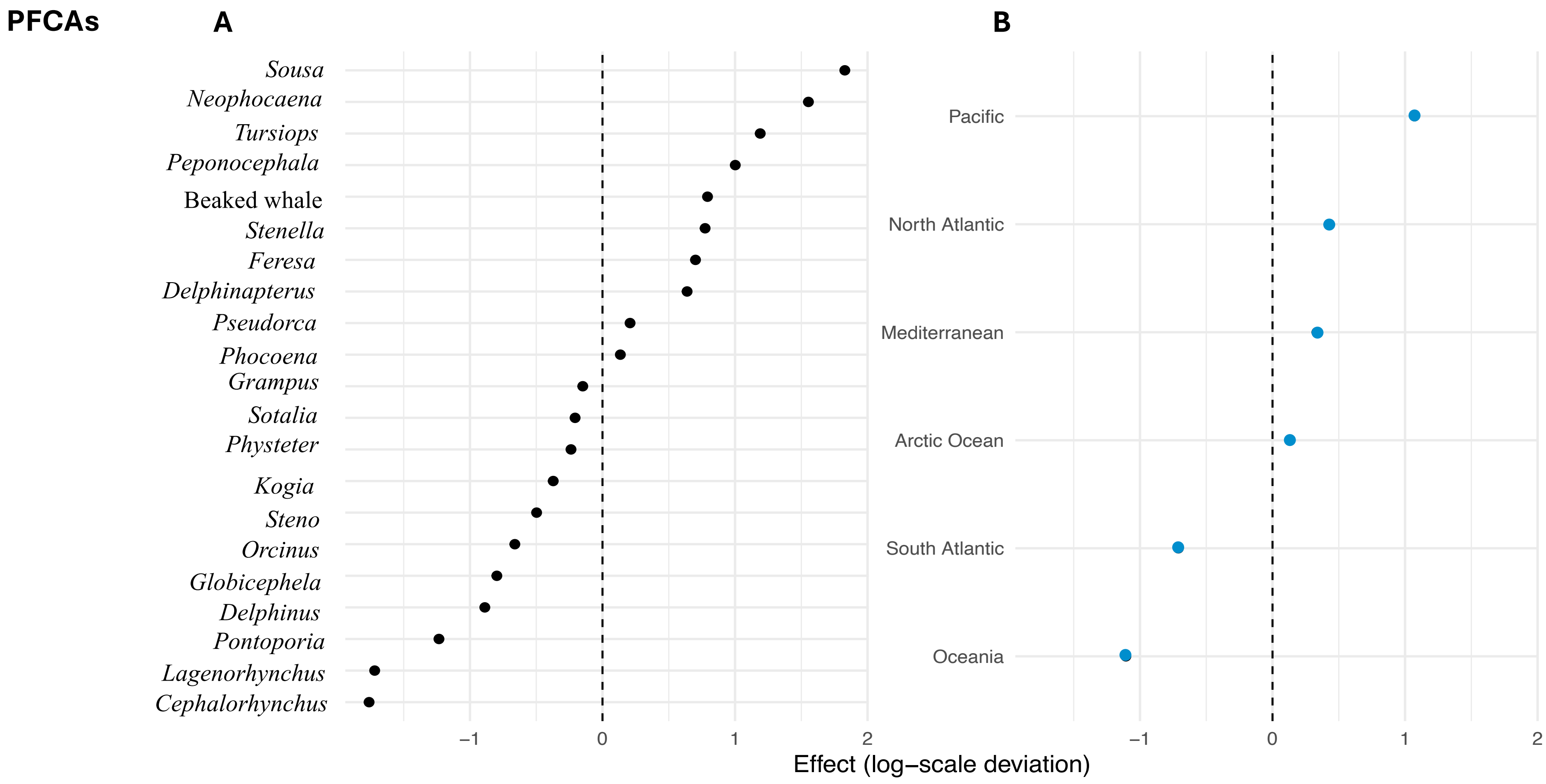

**S16.** Variation in random intercept effects for genus groups A (black) and ocean locations B (blue) from the model predicting log PFOS concentrations. Values represent deviations from the overall intercept on the log scale, with positive values indicating higher contamination and negative values indicating lower contamination relative to the model's grand mean.

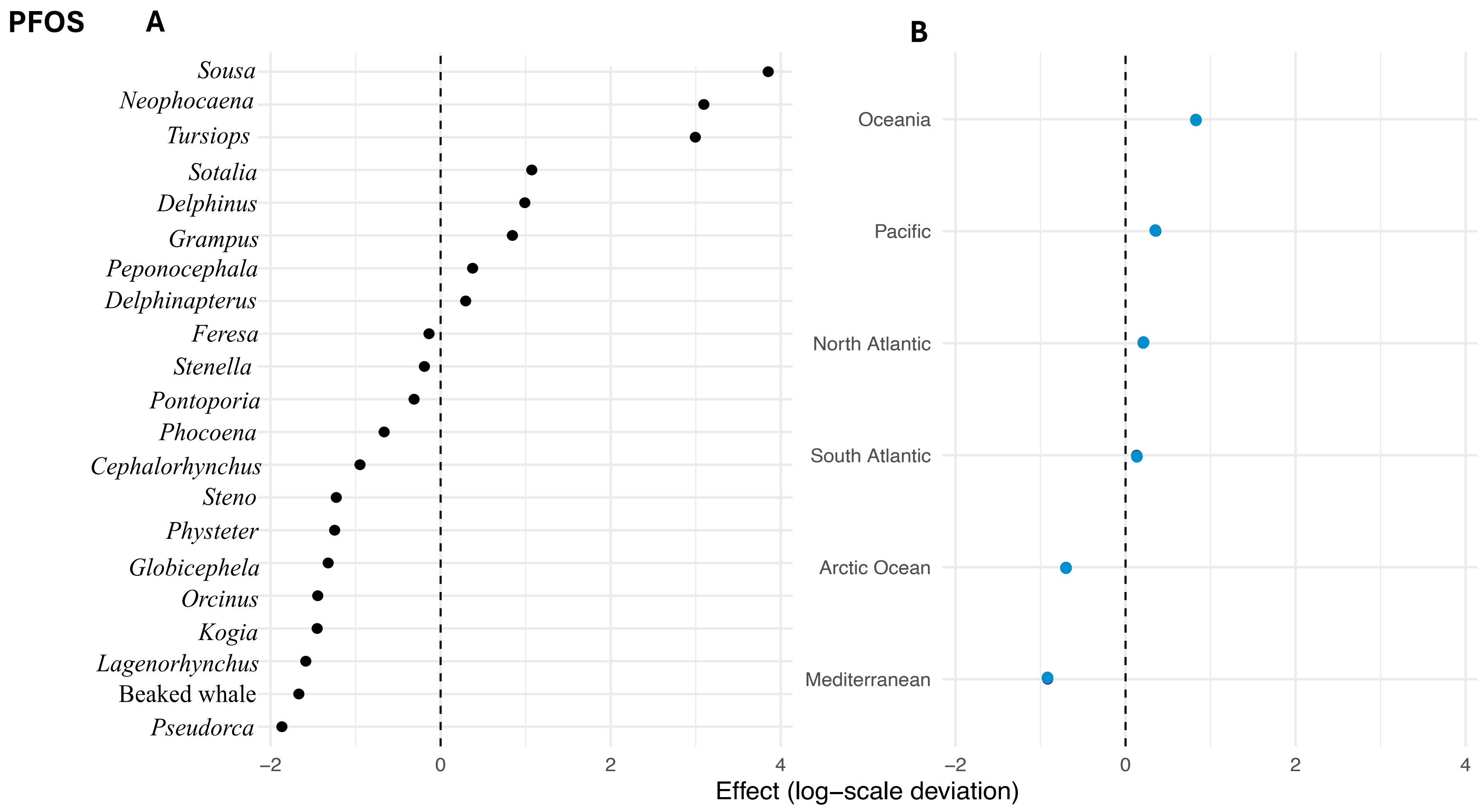

**S17.** Relationship between Analytes and Extracted Internal Standards (EIS) and the Multiple Reaction Monitoring (MRM).

| Analyte (CAS number) | Mass transitions | | Extracted Internal Standard | Type | Mass transition |
|---|---|---|---|---|---|
| PFBA (375-22-4) | 212.9 / 169.0 | | $^{13}C_4$PFBA | Internal | 217.0 / 172.0 |
| PFPeA (2706-90-3) | 262.9 / 219.0 | | $^{13}C_5$PFPeA | Internal | 267.9 / 223.0 |
| PFHxA (307-24-4) | 313.0 / 269.0 | 313.0 / 119.0 | $^{13}C_5$PFHxA | Internal | 318.0 / 273.0 |
| PFHpA (375-85-9) | 363.0 / 319.0 | 363.0 / 169.0 | $^{13}C_4$PFHpA | Internal | 367.0 / 322.0 |
| PFOA (335-67-1) | 413.0 / 369.0 | 413.0 / 169.0 | $^{13}C_8$PFOA | Internal | 421.0 / 376.0 |
| PFNA (375-95-1) | 463.0 / 419.0 | 463.0 / 219.0 | $^{13}C_9$PFNA | Internal | 472.0 / 427.0 |
| PFDA (335-76-2) | 513.0 / 469.0 | 513.0 / 219.0 | $^{13}C_6$PFDA | Internal | 519.0 / 474.0 |
| PFUnDA (2058-94-8) | 563.0 / 519.0 | 563.0 / 169.0 | $^{13}C_7$PFUnDA | Internal | 570.0 / 525.0 |
| PFDoA (307-55-1) | 613.0 / 569.0 | 613.0 / 169.0 | $^{13}C_2$PFDoA | Internal | 615.0 / 570.0 |
| PFTrDA (72629-94-8) | 663.0 / 619.0 | 663.0 / 169.0 | $^{13}C_2$PFDoA | Internal | 615.0 / 570.0 |
| PFTeDA (376-06-7) | 713.0 / 669.0 | 713.0 / 169.0 | $^{13}C_2$PFTeDA | Internal | 715.0 / 670.0 |
| PFPrS (423-41-6) | 249.0 / 80.0 | 249.0 / 99.0 | $^{13}C_3$PFBS | Internal | 302.0 / 99.0 |
| PFBS (375-73-5) | 298.9 / 80.0 | 298.9 / 99.0 | $^{13}C_3$PFBS | Internal | 302.0 / 99.0 |

| | | | | | |
|---|---|---|---|---|---|
| PFPeS (2706-91-4) | 349.0 / 80.0 | 349.0 / 99.0 | $^{13}C_3$PFBS | Internal | 302.0 / 99.0 |
| PFHxS (432-50-7) | 399.0 / 80.0 | 399.0 / 99.0 | $^{13}C_3$PFHxS | Internal | 402.0 / 80.0 |
| PFHpS (375-92-8) | 449.0 / 80.0 | 449.0 / 99.0 | $^{13}C_3$PFHxS | Internal | 402.0 / 80.0 |
| PFOS (1763-23-1) | 499.0 / 80.0 | 499.0 / 99.0 | $^{13}C_8$PFOS | Internal | 507.0 / 99.0 |
| PFNS (474511-07-4) | 549.0 / 80.0 | 549.0 / 99.0 | $^{13}C_8$PFOS | Internal | 507.0 / 99.0 |
| PFDS (335-77-3) | 599.0 / 80.0 | 599.0 / 99.0 | $^{13}C_8$PFOS | Internal | 507.0 / 99.0 |
| PFDoS (79780-39-5) | 699.0 / 80.0 | 699.0 / 99.0 | $^{13}C_8$PFOS | Internal | 507.0 / 99.0 |
| 4:2 FTS (757124-72-4) | 327.0 / 307.0 | 327.0 / 81.0 | $^{13}C_2$4:2 FTS | Internal | 329.0 / 309.0 |
| 6:2 FTS (27619-97-2) | 427.0 / 407.0 | 427.0 / 81.0 | $^{13}C_2$6:2 FTS | Internal | 429.0 / 409.0 |
| 8:2 FTS (39108-34-4) | 527.0 / 507.0 | 527.0 / 81.0 | $^{13}C_2$8:2 FTS | Internal | 529.0 / 509.0 |
| 10:2 FTS (120226-60-0) | 627.0 / 607.0 | 627.0 / 80.0 | $^{13}C_2$8:2 FTS | Internal | 529.0 / 509.0 |
| PFOSA (754-91-6) | 498.0 / 78.0 | 498.0 / 169.0 | $^{13}C_8$PFOSA | Internal | 506.0 / 78.0 |
| N-MeFOSAA (2355-31-9) | 570.0 / 419.0 | 570.0 / 483.0 | $d_3$-N-MeFOSAA | Internal | 573.0 / 419.0 |
| N-EtFOSAA(2991-50-6) | 584.0 / 419.0 | 584.0 / 526.0 | $d_5$-N-EtFOSAA | Internal | 589.0 / 419.0 |
| MeFOSA (31506-32-8) | 512.0 / 169.0 | 512.0 / 219.0 | $d_3$-MeFOSA | Internal | 515.0 / 169.0 |

| | | | | | |
|---|---|---|---|---|---|
| MeFOSE (24448-09-7) | 616.0 / 59.0 | 556.0 / 122.0 | $d_7$-MeFOSE | Internal | 623.0 / 59.0 |
| HFPO-DA (13252-13-6) | 285.0 / 169.0 | 285.0 / 185.0 | $^{13}C_8$HPO-DA | Internal | 287.0 / 169.0 |
| 9Cl-PF3ONS (756426-58-1) | 531.0 / 351.0 | 533.0 / 353.0 | $^{13}C_8$PFOS | Internal | 507.0 / 99.0 |
| 11Cl-PF3OUdS (83329-89-9) | 631.0 / 451.0 | 631.0 / 83.0 | $^{13}C_8$PFOS | Internal | 507.0 / 99.0 |
| ADONA (958445-44-8) | 377.0 / 251.0 | 377.0 / 85.0 | $^{13}C_8$HPO-DA | Internal | 287.0 / 169.0 |
| | | | $^{13}C_3$PFBA | Recovery | 216.0 / 172.0 |
| | | | $^{13}C_2$PFOA | Recovery | 415.0 / 370.0 |
| | | | $^{13}C_2$PFDA | Recovery | 515.0 / 470.0 |
| | | | $^{13}C_4$PFOS | Recovery | 503.0 / 99.0 |
